# Multiscale alterations and cortex-wide organizational trends in focal cortical dysplasia

**DOI:** 10.64898/2026.07.17.739213

**Authors:** Ella Sahlas, Judy Chen, Arielle Dascal, Benjamin Lévesque Kinder, Charlotte Favre, Yigu Zhou, Jessica Royer, Guido I. Guberman, Ke Xie, Jack Lam, Fatemeh Fadaie, Raluca Pana, Jeffery A. Hall, Theodor Rüber, Alexander G. Weil, Andrea Bernasconi, Neda Bernasconi, Boris C. Bernhardt

**Author notes:** Correspondence to: Boris C. Bernhardt, PhD, Montreal Neurological Institute and Hospital (NW-254) 3801 University Street, Montreal, QC H3A 2B4, Canada e, Ella Sahlas, MD-PhD student e.

## Abstract

**BACKGROUND AND OBJECTIVES:** Focal cortical dysplasia (FCD) is a leading cause of surgically remediable pharmaco-resistant epilepsy. Previous research has used various MRI sequences to profile FCD lesions, but multi-site studies using personalized features derived from sequences routinely used in the clinic are sparse. This study aimed to quantify changes in cortical morphology, myeloarchitecture, and function within FCD lesions, compare MRI signatures between lesion subtypes and outcomes, and investigate links to macroscale brain organization.

**METHODS:** This cross-sectional study included patients with FCD-related epilepsy aggregated across 3 datasets from Canada and Germany. The 3T MRI acquisitions included T1-weighted and fluid-attenuated inversion recovery (FLAIR) sequences, with the addition of a resting-state functional sequence for two datasets. We derived cortex-wide maps of patient-specific variations in morphology, myeloarchitecture, and function. Variations were quantified in lesions, ipsilateral cortex, and homotopic regions using age- and sex-adjusted normative models. Subgroup analyses explored the role of histological subtype and surgical outcome. Variations were also related to anterior-posterior and sensory-association organizational gradients.

**RESULTS:** We included 159 patients with epilepsy and FCD (47 % female) and 183 healthy control participants (50 % female). Lesions exhibited changes in morphology (cortical thickness; FLAIR blurring) and in specific measures of depth-dependent intracortical myelin (variance and kurtosis of intracortical myelin profiles across depths), but not in local function (regional homogeneity; node strength). Spatial contextualization indicated more pronounced thickness changes in transmodal association cortices, while increased blurring of the gray-white matter interface co-localized with posterior cortical regions. FCD Type IIb lesions contained more marked changes in blurring and depth-dependent intracortical myelin (kurtosis of intracortical myelin profiles across depths) than Type IIa lesions.

**DISCUSSION:** Our findings provide robust evidence that multiscale MRI profiling can identify FCD signatures and contribute to *in-vivo* subtyping. Furthermore, the novel use of myeloarchitecture profiling and contextualization with macro-scale brain gradients provides new avenues to understand intracortical alterations and the embedding of FCD lesions into broader organizational patterns.

## INTRODUCTION

Focal cortical dysplasia (FCD) is a leading cause of pharmaco-resistant epilepsy, with substantial impacts on health, quality of life, and well-being^1,2^. FCD Type II is a malformation of corticogenesis characterized by cortical dyslamination together with a spectrum of cytological alterations, ranging from the presence of dysmorphic neurons (Type IIa) to the presence of both dysmorphic neurons and balloon cells (Type IIb)^3,4^. FCD Type II occurs earlier in corticogenesis than others malformations of cortical development, such as heterotopia and polymicrogyria^5^, opening a window into the pathophysiology of disruptions in neural proliferation, differentiation, and migration. While FCD Type II is frequently amenable to surgical treatment, both diagnosis and prognosis often represent important clinical challenges. At the diagnostic level, subtle FCD lesions can elude conventional investigations, contributing to delays in establishing surgical candidacy^6^. Notably, previous reports suggest that up to 40% of FCD Type II cases go undiagnosed upon initial imaging^7^. As for prognosis, surgical outcomes vary depending on lesion location and subtype^8^.

The utility of MRI for detecting FCD is recognized clinically, and detailed lesion characterization is critical to treatment planning, from delineation of lesions in surgical candidates to subtyping of lesions, which may inform prognosis^9,10^. Other studies have characterized lesions to evaluate candidate imaging markers^11,12^. Such MRI profiling approaches have highlighted changes in image intensity and diffusion-based metrics in FCD lesions^11,12^. However, previous research has rarely capitalized on normative models and has often been limited by the use of single-site cohorts, hindering generalizability and potential for individualized prediction. Moreover, several studies were based on MRI sequences not included in routine clinical acquisitions, which limits translational potential. We therefore propose a multi-site approach that derives personalized alteration scores from widely used acquisitions, while accounting for patient-specific factors.

Characterization of FCD myeloarchitecture has also been limited to date. Despite the intracortical involvement of FCD lesions, classical MRI markers have included cortical thickness, curvature, and blurring, commonly derived from T1-weighted (T1w) MRI, among other features derived from diffusion-weighted, resting-state functional MRI (rsfMRI), and fluid-attenuated inversion recovery (FLAIR)^13,14^ sequences. Depth-dependent cortical profiling provides markers that may serve as proxies of myeloarchitecture and intracortical lamination, but these features have not yet been applied to FCD^15,16,17^. A prior study conducted microstructural profiling and parametrization of central moments in developing cohorts, obtaining MRI measures of overall and depth-dependent cortical myelin^15^. Another study incorporated intracortical microstructure profiling into a robust workflow operating on both MRI and histological datasets and demonstrated the robustness and scalability of this approach^17^. Capitalizing on these features may help capture dyslamination characteristics of FCD lesions and thereby complement more established features of morphological and intensity change^18,19,20,21^. In addition to identifying changes that are commonly atypical in FCD, the use of multiple features may advance *in-vivo* FCD subtyping^12,10,22^. Indeed, identifying features that could help differentiate subtypes or groups with different surgical outcomes may ultimately aid in prognostication.

Contemporary work has begun to investigate large-scale trends in cortical organization^23^. Of particular relevance have been findings indicating that cortical spatial patterning in the adult brain, as well as neurodevelopmental changes, can be conceptualized and analyzed relative to gradients running in anterior-posterior as well as sensory-transmodal directions. The anterior-posterior gradient has notably been associated with differences in periods of neurogenesis across brain regions, while the sensory-association axis has been shown to capture the progression of developmental plasticity^23,24^. Surprisingly, the profiles of FCD lesions have rarely been related to their location along these gradients. This avenue may help deepen current understandings of how lesion characteristics and their severity may vary as a function of the local milieu in which cortical dyslamination arises. Functional milieus may range, for instance, from lower-order sensory regions to higher-order association cortices. Prior research has related axes of cortical organization to the probability of FCD occurrence^25^, but has not explored links to lesion characteristics. Here, we address these gaps by (*i*) profiling lesions in terms of morphology, myeloarchitecture, and function, (*ii*) investigating the role of histopathological subtype and seizure outcome, and (*iii*) relating gradients to lesional features.

## METHODS

### Participant recruitment

For the Microstructure-Informed Connectomics (MICs) dataset, we recruited patients with a diagnosis of drug-resistant epilepsy due to FCD Type II from the epilepsy monitoring unit of the Montreal Neurological Institute and Hospital between 2023 and 2024. Age- and sex-matched healthy control participants with no history of neurological or psychiatric conditions were recruited from the community between 2021 and 2025. Patient and control participants underwent the same multimodal 3T MRI protocol, and MRI was acquired before surgery in all patients. Patients were selected consecutively based on diagnosis of FCD and availability of both T1w and FLAIR sequences. Diagnosis was determined based on a comprehensive clinical examination including semiology analysis, video-electroencephalogram (EEG) telemetry, clinical neuroimaging, and neuropsychological assessment. Stereo-EEG and histopathology were also examined in patients who were operated. For the Neuroimaging of Epilepsy Laboratory (NOEL) and Bonn datasets, participant recruitment followed similar principles and has been described in detail elsewhere^26,27^.

### MRI data acquisition

MRI data were obtained in patients and control participants, including T1w and FLAIR in all participants^28^ (**Figure 1A**) and rsfMRI in MICs participants and in a subset of NOEL participants (37 of 69 patients and 42 of 48 healthy control participants). Details of data acquisition are described in *Supplemental Material*.

**Figure 1.**
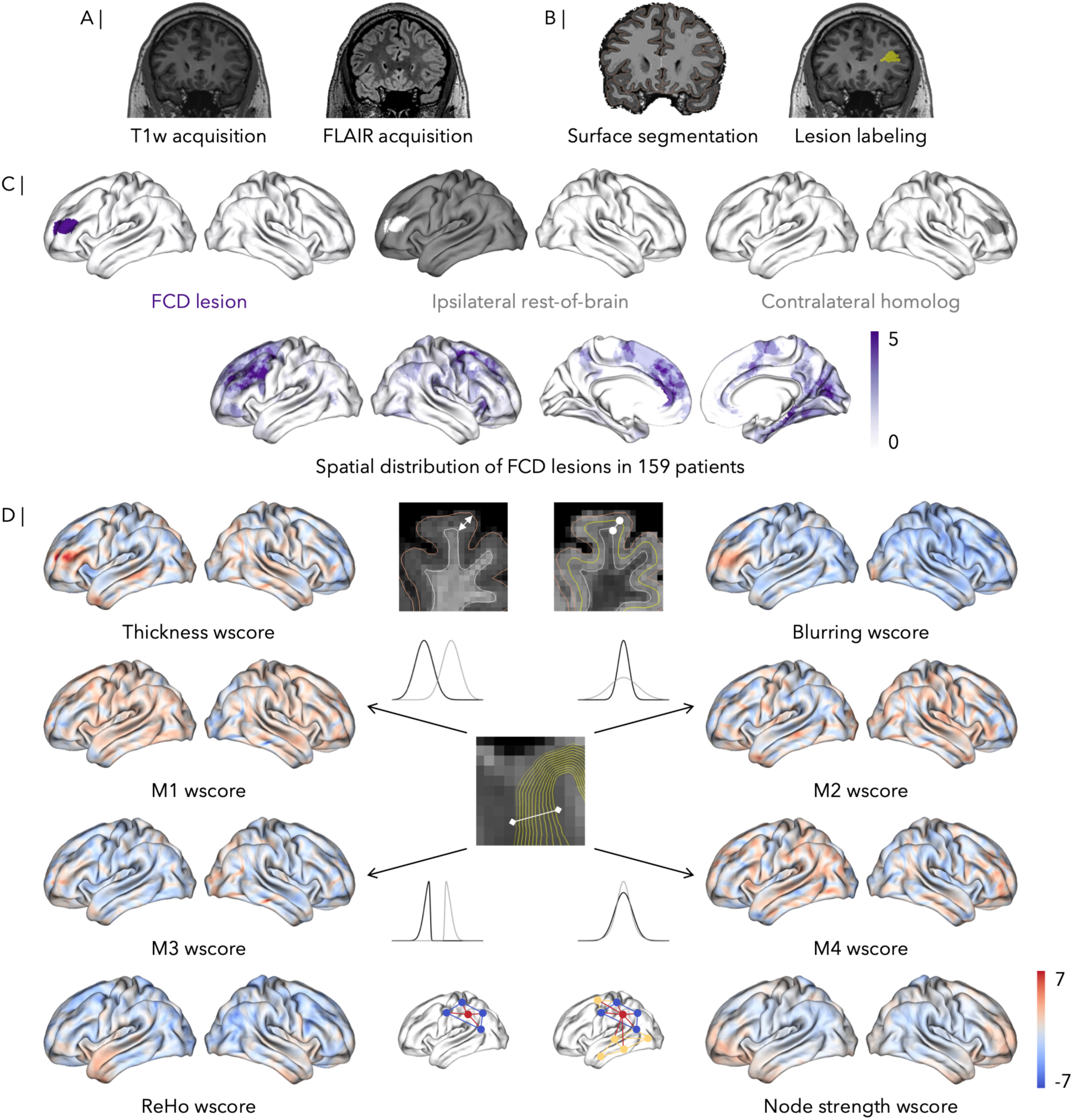
FCD lesion mapping and parallel patient-specific, cortex-wide quantification of changes in cortical morphology, intracortical myeloarchitecture, and function. (a) T1-weighted (T1w) acquisition (*left*); fluid-attenuated inversion recovery (FLAIR) acquisition (*right*). (b) Example of a manually segmented FCD lesion label, segmented in MNI152 space (*left*); example of FreeSurfer segmentations of the pial matter and white matter surfaces, with manual quality control for each case (*right*). (c) Projection of FCD lesion labels to a standardized surface space in a representative patient (*top, left*). Identification of all ipsilateral cortical locations outside of the lesion label (*top, middle*). Identification of the cortical locations mirroring the lesion label in the contralateral hemisphere, or homotopic region (*top, right*). Map of the distribution of FCD lesion locations across the cortex for all 159 patients (*bottom*). (d) Cortex-wide maps of patient-specific changes in cortical thickness, FLAIR blurring, M1 (mean of intracortical myelin across cortical depths), M2 (variance across depths), M3 (skewness across depths), M4 (kurtosis across depths), regional homogeneity, and node strength, computed as the *w*-score relative to matched control cohorts at each cortical location.

### MRI data processing

We processed 3T MRI data from all sites using Micapipe (version 0.2.3, https://micapipe.readthedocs.io)^29^, a robust open-access pipeline allowing for the unified preprocessing of multimodal MRI data, to extract surface-based measures of cortical microstructure and function.

T1w scans were reoriented to standard orientation, deobliqued, corrected for intensity nonuniformity, intensity normalized, and skull stripped. We generated cortical surface models from native T1w scans for each participant using FreeSurfer (version 6.0, https://surfer.nmr.mgh.harvard.edu)^30^, an open-source toolbox allowing for the delineation of the gray and white matter surfaces (**Figure 1B**). For all participants, rigorous manual FreeSurfer quality control was conducted to correct for segmentation errors in surface extraction via placement of control points and manual edits.

FLAIR images underwent gray matter and white matter segmentation, bias field correction weighted by white matter, normalization of intensities by the peak (or mode) of the white matter tissue intensities, brain masking, and registration to the individual native processing space.

Vertex-wise native maps of cortical thickness (CT) were generated in native surface space for each participant by computing the Euclidian distance separating corresponding pial and white matter vertices. Vertex-wise maps of FLAIR intensity were also generated in native surface space.

Using Connectome Workbench^31^, an open-source tool for mapping neuroimaging data, we registered native cortical features—CT, FLAIR intensity, and rsfMRI timeseries—to the fsLR-32k template surface, with 32492 vertices per hemisphere.

### Structural feature generation

We computed FLAIR blurring (BL) at each vertex as the difference between white matter FLAIR intensity and mid-thickness FLAIR intensity, divided by the distance between the white matter and mid-thickness surfaces (i.e. half of the thickness of the cortex at a given vertex).

We applied intensity non-uniformity correction^29^ to the FLAIR images and intensity clamping, rescaling, and normalization to both the FLAIR and T1w images. We then computed the T1w/FLAIR ratio, which provides a contrast that can be used to derive myeloarchitectural markers of interest for datasets in which a microstructure-sensitive contrast was not directly acquired^32^. We sampled intracortical intensities along 12 equivolumetric surfaces between the pial and white matter boundaries. We used the resulting intensity profiles to compute the first (M1), second (M2), third (M3), and fourth (M4) moments, or mean, variance, skewness, and kurtosis of intensities across the 12 cortical depths, respectively. These measures of myeloarchitecture can be interpreted as proxies of cortical myelin content and shifts in myelin content across cortical depths.

We applied spatial smoothing (Gaussian kernel, full width at half maximum (FWHM) = 10 mm) to the CT, FLAIR intensity (sampled at both the white matter and mid-thickness surfaces), M1, M2, M3, and M4 maps.

### Functional feature generation

From rsfMRI timeseries, we derived maps of regional homogeneity (RH) and node strength (NS). RH, a proxy of regional connectivity, was computed as the temporal coherence of rsfMRI timeseries among the nearest regional neighbours of a given vertex^33,34^. NS, a proxy of global connectivity, was computed as the sum of the weights of all functional connections for a given vertex after generating a functional connectivity matrix comprising Pearson correlation coefficients between the rsfMRI timeseries for each vertex^33,35^. We applied spatial smoothing (Gaussian kernel, FWHM = 10 mm) to the RH and NS maps.

### Feature harmonization

We applied ComBat harmonization^36^ to the CT, BL, M1, M2, M3, M4, RH, and NS data to control for the confounding effect of site (or scanner) across all datasets while preserving effects of age, sex, and group, as in prior work^37^. The NOEL dataset included patients scanned using two different scanners over time and was therefore treated as two datasets for harmonization.

### Lesion labels

Lesions of patients in the MICs dataset were manually labeled by E.S. under the supervision of two epilepsy imaging experts (A.B. and N.B.). A.B., N.B., G.G., and E.S. contributed to reaching a consensus on lesion borders via discussion. Lesion segmentation was completed before quantitative MRI feature generation. Segmentation was performed using both T1w and FLAIR images co-registered to MNI152 space, with intensity non-uniformity correction and clamping of intensities between 0 and 100. Lesions of patients in the NOEL dataset were labeled by A.B. and N.B. using the same procedures (**Figure 1B**). Lesions of patients in the Bonn dataset were similarly manually labeled via the collaboration of two neurologists experienced in epilepsy imaging, as described in Schuch *et al.*^27^.

For each patient, we mapped the volumetric lesion mask to the native surface, then proceeded with resampling to the fsLR-32k template surface (**Figure 1C**). In addition, we located the ipsilateral vertices outside of the lesion mask, and we located the vertices mirroring the lesion mask in the contralateral hemisphere, or homotopic region (**Figure 1C**).

### Mapping changes in microstructure, myeloarchitecture, and function

We computed vertex-wise *w*-scores quantifying alterations in CT, BL, M1, M2, M3, M4, RH, and NS in each patient relative to the corresponding control cohort while adjusting for age and sex (**Figure 1D**). These *w*-scores can be interpreted similarly to *z*-scores, capturing deviations from the norm in each feature. The following formula was used to estimate *w*-scores:

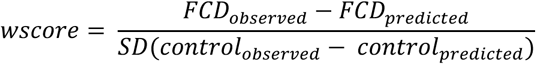

### Quantifying alterations in lesions, ipsilateral non-lesional cortex, and homotopic regions

For each feature, we compared mean *w*-scores within patient-specific lesions to mean *w*-scores across the rest of the ipsilateral cortex. As an additional control comparison, we compared mean *w*-scores within lesions to mean *w*-scores within the homotopic region. Comparisons were performed using paired-samples *t*-tests, and we controlled for multiple comparisons using False Discovery Rate (FDR) correction.

### Subgroup analyses

We compared lesional *w*-scores profiles for CT, BL, M1, M2, M3, M4, RH, and NS between subgroups defined by histopathological subtype of FCD lesions and surgical outcomes for operated patients. Specifically, mean within-lesion *w*-scores for each feature were compared between (1) patients with FCD Type IIa versus Type IIb and (2) patients with Engel I versus non-Engel I outcomes following surgery. We used paired-samples *t*-tests, and FDR corrections were applied to control for multiple comparisons.

### Gradient associations

To investigate the potential associations between features showing significant alterations within lesions and gradients of brain organization, we correlated the difference between mean within-lesion and mean ipsilateral non-lesional cortex *w*-scores for CT, BL, M2, and M4 with mean within-lesion rank along anterior-posterior and sensory-association functional connectivity gradients. We computed Pearson correlations, and FDR corrections were applied to control for multiple comparisons. We used the functional connectivity gradient provided by BrainSpace^38^, which is computed from the HCP dataset^39,40^.

### Standard protocol approvals, registrations, and patient consents

*Research ethics board approval.* The Research Ethics Board of the Montreal Neurological Institute and Hospital approved this research involving human participants. *Informed consent.* All participants provided written informed consent.

### Data availability

Preprocessed MRI feature maps used in the study will be made available on the Open Science Framework.

## RESULTS

### Participants

We studied 159 patients with epilepsy and FCD Type II (74 women and 85 men) and 183 age-matched healthy control participants (91 women and 92 men) across 3 sites located in Montreal, Canada (MICs and NOEL datasets) and Bonn, Germany (Bonn dataset). The MICs dataset included 5 FCD patients (3 women and 2 men, with a mean age ± SD of 26.8 ± 5.5 years) and 50 control participants (25 women and 25 men, 27.5 ± 4.5 years). The NOEL dataset included 69 FCD patients (36 women and 33 men, 24.6 ± 9.8 years) and 48 control participants (23 women and 25 men, 31.7 ± 8.0 years). The Bonn dataset included 85 FCD patients (35 women and 50 men, 28.9 ± 12.4 years) and 85 control participants (43 women and 42 men, 33.3 ± 11.9 years). These data are summarized in **Table 1**. Among patients with histopathological data available after surgery (N = 115/159), 51 had FCD Type IIa and 64 had FCD Type IIb. Finally, among patients with seizure outcome data available following surgery (N = 114/159), 85 had seizure-free (SF, Engel I) outcomes and 29 had non-seizure-fee (NSF, Engel II-IV) outcomes.

**Table 1.** Participant demographics.

|  | **MICs dataset** | **NOEL dataset** | **Bonn dataset** |
| --- | --- | --- | --- |
| **Number** | 5 patients<br>50 controls | 69 patients<br>48 controls | 85 patients<br>85 controls |
| **Sex (F/M)** | 3/2<br>25/25 | 36/33<br>23/25 | 35/50<br>43/42 |
| **Age at scan** (years ± SD) | 26.8 ± 5.5<br>27.5 ± 4.5 | 24.6 ± 9.8<br>31.7 ± 8.0 | 28.9 ± 12.4<br>33.3 ± 11.9 |

### Alterations in candidate markers within FCD lesions relative to ipsilateral cortex

Group-wise analyses revealed increased *w*-scores within patient-specific lesions compared to the rest of the ipsilateral cortex for CT (mean ± SE: 0.971 ± 0.139 *vs* 0.023 ± 0.041; paired *t*-test: *t* = 7.151, *p*_FDR_ = 1.609*10^-10^; **Figure 2A**), BL (2.220 ± 0.146 *vs* 0.951 ± 0.127; *t* = 13.577, *p*_FDR_ = 4.073*10^-27^; **Figure 2A**), and M4 (0.403 ± 0.065 *vs* 0.120 ± 0.042; *t* = 4.748, *p*_FDR_ = 1.046*10^-5^; **Figure 2B**). Group-wise analyses also revealed decreased *w*-scores within patient-specific lesions compared to the rest of the ipsilateral cortex for M2 (-0.787 ± 0.079 *vs* -0.321 ± 0.062; *t* = -6.714, *p*_FDR_ = 1.289*10^-9^; **Figure 2B**) and NS (-0.294 ± 0.192 *vs* 0.040 ± 0.167; *t* = -2.864, *p*_FDR_ = 1.166*10^-2^; **Figure 2C**). However, there were no significant differences between *w*-scores within lesions and across the rest of the ipsilateral cortex for M1 (-0.495 ± 0.094 *vs* -0.393 ± 0.065; *t* = -1.563, *p*_FDR_ = 1.748*10^-1^; **Figure 2B**), M3 (0.482 ± 0.094 *vs* 0.391 ± 0.065; *t* = 1.377, *p*_FDR_ = 2.272*10^-1^; **Figure 2B**), and RH (-0.156 ± 0.124 *vs* -0.077 ± 0.063; *t* = -0.655, *p*_FDR_ = 5.581*10^-1^; **Figure 2C**).

**Figure 2.**
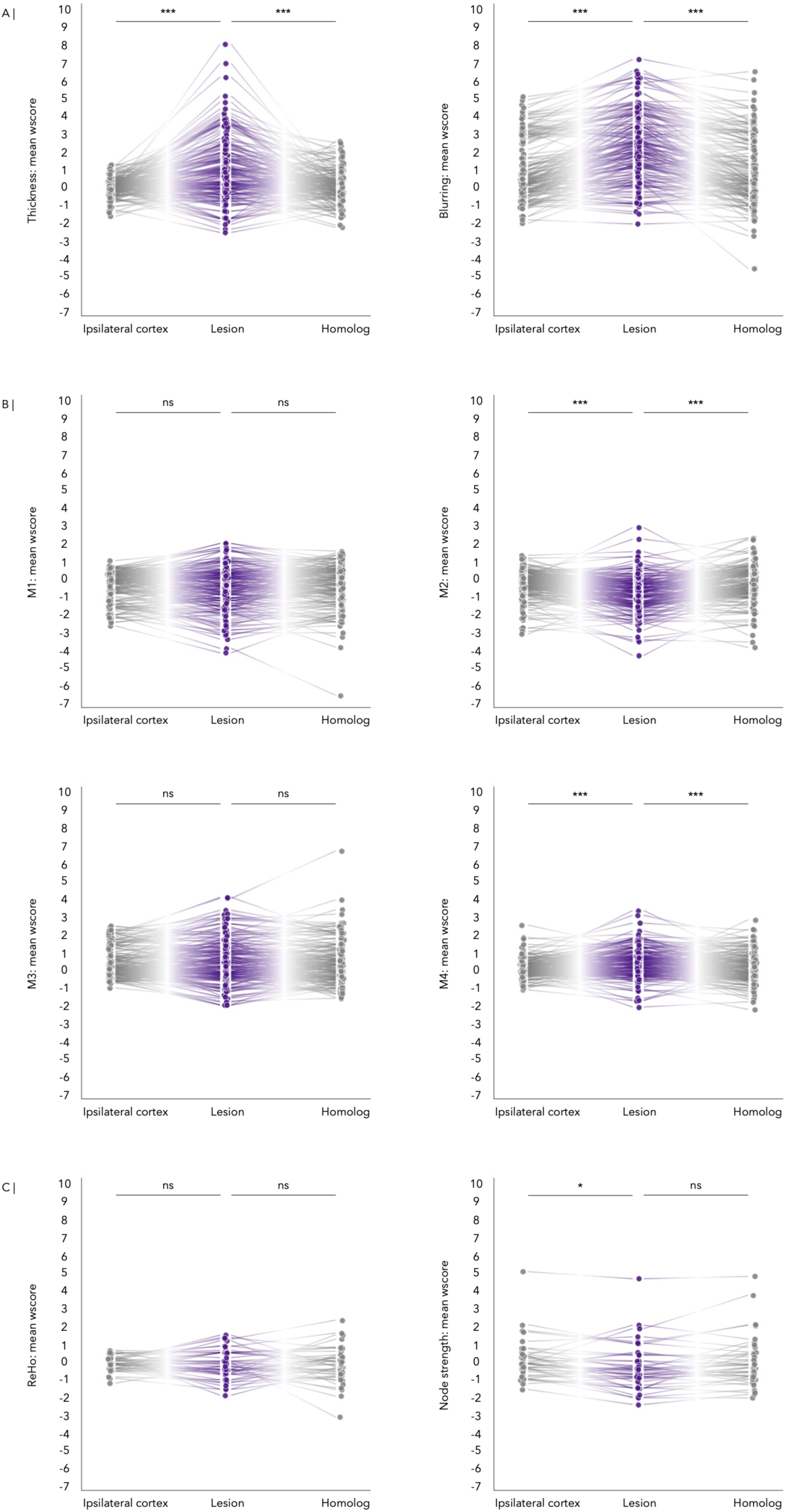
Multimodal profiling of FCD and non-lesional cortex. (**a**) Comparison of mean cortical thickness *w*-scores within patient-specific lesions to mean scores across ipsilateral cortex and in homotopic regions (*left*). Comparison of mean FLAIR blurring *w*-scores within patient-specific lesions to mean scores across ipsilateral cortex and in homotopic regions (*right*). (**b**) Comparison of mean M1 *w*-scores within patient-specific lesions to mean scores across ipsilateral cortex and in homotopic regions (*top, left*). Comparison of mean M2 *w*-scores within patient-specific lesions to mean scores across ipsilateral cortex and in homotopic regions (*top, right*). Comparison of mean M3 *w*-scores within patient-specific lesions to mean scores across ipsilateral cortex and in homotopic regions (*bottom, left*). Comparison of mean M4 *w*-scores within patient-specific lesions to mean scores across ipsilateral cortex and in homotopic regions (*bottom, right*). (**c**) Comparison of mean regional homogeneity *w*-scores within patient-specific lesions to mean scores across ipsilateral cortex and in homotopic regions (*left*). Comparison of mean node strength *w*-scores within patient-specific lesions to mean *w*-scores across ipsilateral cortex and in homotopic regions (*right*). Across all panels: ***: *p*_FDR_ < 0.001; *: *p*_FDR_ < 0.05; ns: not significant.

### Alterations in candidate markers within FCD lesions relative to homotopic regions

Group-wise analyses revealed increased *w*-scores within patient-specific lesions compared to homotopic regions for CT (0.971 ± 0.139 *vs* 0.068 ± 0.072; *t* = 6.566, *p*_FDR_ = 2.258*10^-9^; **Figure 2A**), BL (2.220 ± 0.146 *vs* 0.914 ± 0.142; *t* = 13.030, *p*_FDR_ = 6.425*10^-26^; **Figure 2A**), and M4 (0.403 ± 0.065 *vs.* 0.148 ± 0.065; *t* = 3.794, *p*_FDR_ = 4.220*10^-4^; **Figure 2B**). We observed decreased *w*-scores within patient-specific lesions compared to homotopic regions for M2 (-0.787 ± 0.079 *vs* -0.342 ± 0.079; *t* = -5.993, *p*_FDR_ = 3.583*10^-8^; **Figure 2B**). However, there were no significant differences between *w*-scores within lesions and homotopic regions for M1 (-0.495 ± 0.094 *vs* -0.428 ± 0.091; *t* = -0.978, *p*_FDR_ = 4.053*10^-1^; **Figure 2B**), M3 (0.482 ± 0.094 *vs* 0.438 ± 0.093; *t* = 0.640, *p*_FDR_ = 5.581*10^-1^; **Figure 2B**), RH (-0.156 ± 0.124 *vs* -0.151 ± 0.149; *t* = -0.029, *p*_FDR_ = 9.771*10^-1^; **Figure 2C**), and NS (-0.294 ± 0.192 *vs* -0.047 ± 0.202; *t* = -2.196, *p*_FDR_ = 5.411*10^-2^; **Figure 2C**).

### Comparisons across lesion locations, histopathological subtypes, and surgical outcomes

Subgroup analyses comparing mean *w*-scores within Type IIb lesions compared to Type IIa lesions revealed increased BL (2.615 ± 0.224 *vs* 1.682 ± 0.216; *t* = -2.945, *p*_FDR_ = 3.142*10^-2^; **Figure 3B**) and M4 (0.504 ± 0.097 *vs* 0.130 ± 0.106; *t* = -2.597, *p*_FDR_ = 4.263*10^-2^; **Figure 3B**) in the former.

**Figure 3.**
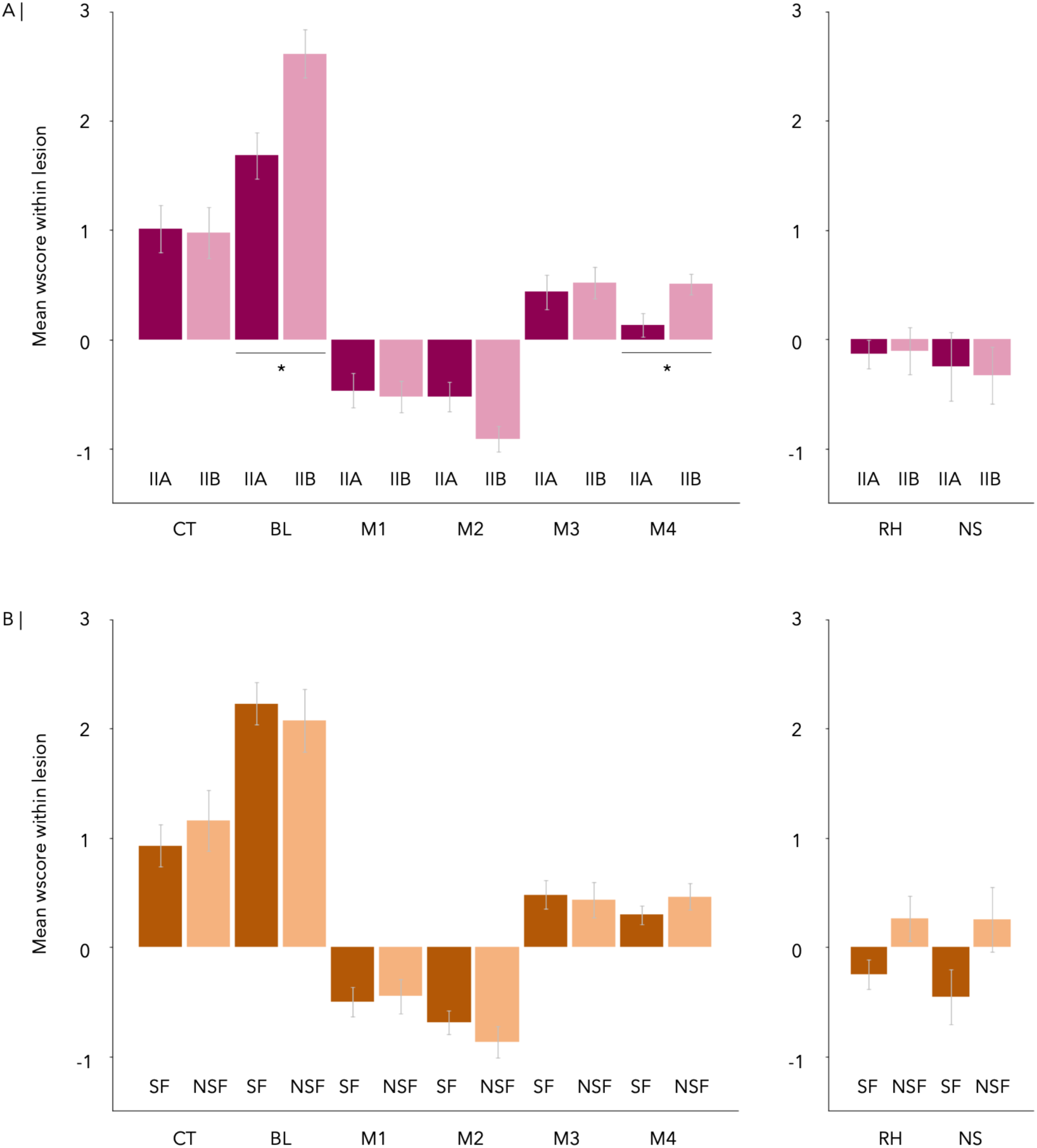
Subgroup analyses based on anatomical, histopathological, and clinical characteristics. (**a**) Comparisons between FCD Type IIa and Type IIb lesions for CT, BL, M1, M2, M3, M4 (*left*) and for RH and NS (*right*). *: *p*_FDR_ < 0.05; other comparisons were not significant. (**c**) Comparisons between FCD lesions of patients with seizure-free (SF, Engel I) post-surgical outcomes and those of patients with non-seizure-free (NSF, Engel II-IV) post-surgical outcomes. No comparisons were significant following FDR corrections.

Comparing mean *w*-scores in patients with seizure-free versus non-seizure-free outcomes did not reach significance for CT, BL, M1, M2, M3, M4, RH, and NS (*p*_FDR_ > 0.05; **Figure 3C**), although the directionality of *w*-scores was opposite in patients with seizure-free outcomes compared to those with non-seizure-free outcomes for RH (-0.248 ± 0.138 *vs* 0.263 ± 0.220; *t* = -1.872, *p*_FDR_ = 5.563*10^-1^; **Figure 3C**) and NS (-0.458 ± 0.257 *vs* 0.252 ± 0.316; *t* = -1.452, *p*_FDR_ = 6.214*10^-1^; **Figure 3C**).

### Relationships between MRI profiles and gradients of brain organization

To elucidate potential links between the MRI signature of a lesion and its location of along key axes of neurodevelopment, we investigated the relationships between each of the four informative features (those that showed significant differences within lesions relative to ipsilateral cortex and homotopic regions) and two main gradients of brain organization. Specifically, we focused on the difference between mean within-lesion *w*-scores and mean ipsilateral cortex *w*-scores for CT, BL, M2, and M4, and on mean within-lesion rank along anterior-posterior and sensory-association functional connectivity gradients.

Following FDR corrections for multiple comparisons, two significant associations emerged from these alteration-gradient correlations. Firstly, there was a negative correlation between the BL difference and the anterior-posterior axis (*r* = -0.209, *p*_FDR_ = 3.312*10^-2^; **Figure 4A**). This finding suggests that lesions located more posteriorly may exhibit a greater extent of blurring relative to non-lesional cortex compared to lesions located more anteriorly. Secondly, there was a positive correlation between the CT difference and the sensory-association hierarchy (*r* = 0.234, *p*_FDR_ = 2.419*10^-2^; **Figure 4B**). This indicates a tendency for lesions located in transmodal association cortices to show more pronounced thickening relative to non-lesional cortex compared to lesions located in unimodal sensorimotor cortices.

**Figure 4.**
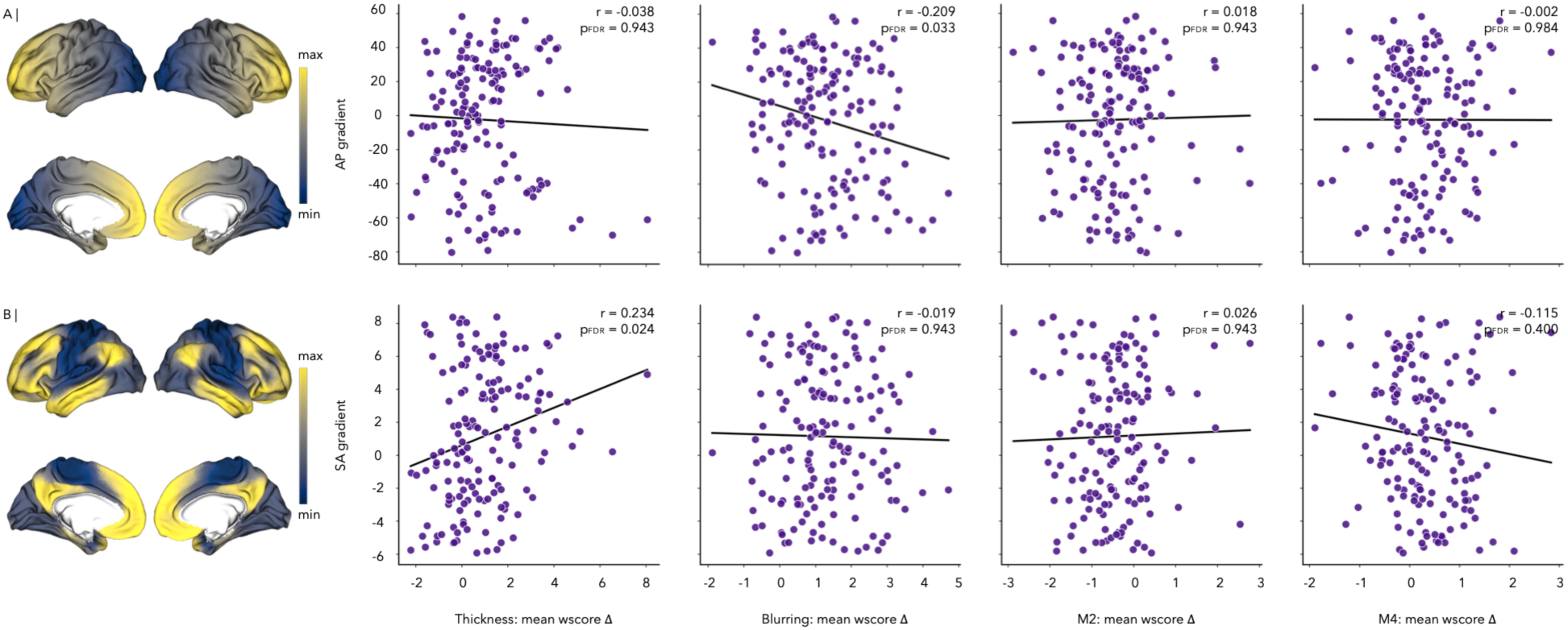
Links between lesional alteration burden and gradients of brain organization. (**a**) Correlations between mean rank along the (**a**) anterior-posterior gradient and (**b**) functional gradient and CT, FL, M2, and M4 alteration burden within lesions across patients. Each dot represents a patient.

## DISCUSSION

Harnessing quantification of widely used MRI sequences and a multi-site, personalized approach, our study suggests that FCD lesions (*i*) contain thickness alterations together with alterations in FLAIR blurring and depth-dependent cortical myelin, (*ii*) exhibit profiles that vary with subtype, and (*iii*) differ depending on their location along two gradients of brain organization. These insights were facilitated by combining tools from the fields of neuroanatomy and neurodevelopment—intracortical microstructure mapping^15,16,17^ and gradient analysis^23,24^—with an approach to quantifying the characteristics of FCD and its subtypes that is compatible with clinical neuroimaging. Findings reveal differences in specific classical and novel candidate markers within FCD lesions relative to non-lesional tissue, both across the ipsilateral cortex and in homotopic regions. Overall, results suggest that personalized multimodal MRI quantification may capture FCD signatures *in-vivo*. Mapping epicentres of alterations in morphology and myeloarchitecture using a *w*-score approach may potentially help enhance techniques aiming to identify and classify epilepsy-causing FCD lesions, which could ultimately contribute to targeted diagnosis, prognosis, and treatment planning. Results suggest that measures of functional coherence and connectivity may not show marked alterations within FCD lesions, warranting further studies focused on identifying MRI-sensitive functional hallmarks of FCD. Moreover, the present study indicates that MRI profiles of FCD Type IIa and IIb may differ not only in terms of blurring alterations, but also in terms of intracortical myelin content across cortical depths. Taken together, the application of intracortical myeloarchitecture and gradient mapping methods to the study of FCD reveals potential to capture both micro-scale dyslamination and macro-scale embeddings.

Findings are consistent with current insights into FCD cytoarchitecture from the field of neurobiology, from meso-scale disruption of the gray-white matter interface to micro-scale changes in intracortical lamination. An immunohistochemistry study of layer-specific cortical neuron subtypes and composition in FCD suggested that while neuronal diversity and general layering may be preserved in FCD Type IIb, losses and increases in specific neuron populations may selectively affect the deepest layers of the cortex^41^. This notion is consistent with our findings regarding myeloarchitecture, whereby the mean intracortical myelin content is preserved within FCD lesions, whereas we observe alterations in two depth-dependent measures of shifts in intracortical myelin across cortical layers within lesions. Consistent with a recent study introducing microstructure profiling as a promising technique to index cortical lamination across various modalities^17^, our application of this technique to the study of FCD lesions yields biologically and clinically plausible results. Furthermore, the finding of increased alterations in FLAIR blurring and depth-dependent variance in intracortical myelin within Type IIb relative to IIa lesions is also consistent with the neurobiological literature. Specifically, histopathological definitions of FCD subtypes indicate that Type IIb is differentiable from IIa by the presence of balloon cells, which are hypertrophic neurons pathognomonic of malformations of cortical development and characterized by a cytoplasmic accumulation of neurofilament proteins that may occur in specific layers^42,43^. The occurrence of such cells could contribute to increased alterations both in FLAIR blurring and in intracortical myelin content variance across layers.

In this study, we focus on profiling and characterization of FCD lesions, with the objective that future studies in independent cohorts may build upon these findings both to expand existing work on MRI-based lesion profiling and to develop or enhance artificial intelligence and machine learning models supporting FCD detection, localization, classification, and prognostication^6,18,19,20,44,45^. One insight applicable to such studies that emerges from the present work is that a neurobiologically grounded approach to feature engineering may carry benefits in terms of lesion detection. As such, we observed the largest differences between lesions and non-lesional ipsilateral cortex for FLAIR blurring, followed by thickness, intracortical myelin variance, and intracortical myelin kurtosis, which, together, were the four most informative features in our study. Functional coherence and connectivity could not detect such differences in the current study, which could be explained by more global epilepsy-related shifts in function than in structural measures, or by the functional heterogeneity of FCD lesions^22^. Meanwhile, although blurring is traditionally derived from T1w MRI^46^, our study highlights FLAIR-derived blurring as a quantitative feature that may supersede thickness in terms of within-lesion feature changes. A second insight lies in the potential relevance of indexing cortical myeloarchitecture for profiling and subtyping of FCD lesions, which, to our knowledge, has yet to be applied in studies of malformations of cortical development. A third insight lies in the relevance and feasibility of deriving candidate markers from sequences that are part of the Harmonized Neuroimaging of Epilepsy Structural Sequences (HARNESS-MRI) protocol^28^. Having increasingly gained adoption in recent years^47^, this protocol is endorsed by the Neuroimaging Task Force of the International League Against Epilepsy with the aim to optimize the use of MRI to benefit patients undergoing clinical evaluations for epilepsy worldwide. As the HARNESS-MRI protocol becomes adopted worldwide, deriving features from its constituent sequences will become increasingly feasible across sites, facilitating future translation.

We further expand FCD characterization to the investigation of associations between the relative magnitude of patient-specific lesional changes and the position of lesions along two axes of brain organization, namely the anterior-posterior and sensory-association functional connectivity gradients. These axes are closely tied to corticogenesis and neurodevelopment, including cortical differentiation, arealization, and myelination processes, and they may also reflect spatial and temporal variations in developmental plasticity and environmental susceptibility across different brain regions^24^. We utilize a functional gradient derived from independent data from the Human Connectome Project^39,40^. Together with open-access processing tools allowing for rapid, standardized registration between observations across modalities, such as Micapipe^29^ and BrainSpace^38^, the availability of such datasets opens the way for previously underexplored avenues of research into the relationship between the macro-scale embedding and the micro-scale signatures of FCD and other malformations of cortical development. Our findings suggest that lesions located more posteriorly may carry more marked FLAIR blurring changes, and lesions located higher in the sensory-association hierarchy may show greater relative thickening. The accentuation of blurring changes in more posterior regions may be explained in part by the high myelin content of some of these regions, such as the striate cortex. Nevertheless, we cannot exclude a role for selection bias, whereby posterior FCD lesions may be overlooked more frequently than typical frontal lesions, unless they are more markedly blurred. In addition, posterior lesions in the visual cortex can be deemed inoperable and patients may not be referred. The greater thickness changes in lesions in heteromodal association cortex could be partially explained by the greater thickness of these areas relative to sensory cortices. Additional research linking other gradients^48,49^ to FCD characteristics may further shed light on the embedding of FCD lesions into broader organizational patterns. Prior work notably indicated that the embedding of lesions into different functional networks is linked to age of epilepsy onset, highlighting the interrelation between brain organization and disease characteristics^50^.

The present work has limitations, highlighting the need for additional multi-site, quantitative studies that may help bridge the remaining gaps in the discovery, detailed characterization, and clinical translation of MRI markers of FCD lesions. First, while both the overall patient cohort (N = 159) and the control cohort (N = 183) are sizeable compared to previous studies in the field^11,12,14^, subgroup analyses inevitably compared more modest groups. This is most notable for the subgroup analysis comparing lesion characteristics between different patients with surgical outcomes, as not all patients underwent surgery at each site, and not all patients who underwent surgery had follow-up and seizure outcome data available. In the future, identifying differences in lesion characteristics associated with different post-operative outcomes will be critical to working towards quantitative tools that can support prognostication. Larger cohort sizes will also facilitate the discovery and thorough evaluation of diagnostic markers. Second, this study attempts to balance detailed anatomical profiling with translational potential by focusing primarily on structural features derived from T1w and FLAIR, sequences that are widely used as part of comprehensive clinical epilepsy evaluations. Nonetheless, exploring features derived from additional sequences that may only be available in research settings may provide more comprehensive lesion profiling. Third, the present analyses remain group-level comparisons of personalized markers, with future potential for further individualization.

The present work examined FCD signatures across and between lesion locations, subtypes, and outcomes, combining classical and novel imaging markers and studying links between lesion signatures and gradients. Patient cohorts span 3 sites located in Canada and Germany, helping to improve the generalizability of findings, and key features are derived from sequences commonly acquired in the clinical setting. Findings are consistent with and build upon current understandings of the neuroanatomical and neurobiological underpinnings of FCD and provide new insights into lesional myeloarchitecture. These results may inform future work aiming to develop new surface-based machine learning models or enhance existing models for localization and classification of FCD lesions. Assessment of the correspondence between FCD lesion characteristics and gradients of brain organization reveals that blurring and thickening changes may relate to anterior-posterior and functional axes, respectively. Additional studies may employ the present framework to investigate other candidate markers of FCD, retain the most informative, and build more accurate and generalizable detection models, with an ultimate goal of perfecting models also capable of aiding in prognostication based on quantitative lesion characteristics, supporting personalized epilepsy care.

## ACKNOWLEDGEMENTS

E.S. and J.C. acknowledge funding from the Canadian Institutes of Health Research (CIHR) Vanier Canada Graduate Scholarships. K.X. acknowledges funding from the Savoy Foundation. J.L. is funded by the Fonds de Recherche du Québec – Santé (FRQS) and the Ministère de la Santé et des Services sociaux du Québec (MSSS). T.R. is funded by the German Federal Ministry of Research, Technology and Space (epi-center.ai) and received payments for lectures from Angelini, Desitin, Eisai, Jazz Pharmaceuticals, and LivaNova. B.C.B. acknowledges funding from CIHR (FDN-154298, PJT-174995), the Natural Sciences and Engineering Research Council of Canada (NSERC) (Discovery-1304413), FRQS, the Brain Canada Foundation, the SickKids Foundation (NI17-039), the Azrieli Centre for Autism Research (ACAR), the Helmholtz International BigBrain Analytics and Learning Laboratory (HIBALL), Healthy Brains, Healthy Lives (HBHL), the Canada Research Chairs Program, and the Centre of Excellence in Epilepsy at the Montreal Neurological Institute-Hospital.

## AUTHOR CONTRIBUTIONS

E.S. and B.C.B. contributed to conception and design of the study; E.S., J.C., A.D., J.R., T.R., and B.C.B. contributed to data acquisition; E.S. contributed to processing of MRI data; E.S., J.C., A.D., B.L., C.F., Y.Z., and J.R. contributed to quality control of MRI data; E.S. and B.C.B. analyzed the data and wrote the manuscript; all authors revised and approved the manuscript.

## REFERENCES

1. Cohen NT, Chang P, You X, et al. Prevalence and risk factors for pharmacoresistance in children with focal cortical dysplasia-related epilepsy. Neurology. 2022;99(18):e2006–e2013. doi:10.1212/WNL.0000000000201033

2. Diaz RJ, Sherman EM, Hader WJ. Surgical treatment of intractable epilepsy associated with focal cortical dysplasia. Neurosurg Focus. 2008;25(3):E6. doi:10.3171/FOC/2008/25/9/E6

3. Blümcke I, Thom M, Aronica E, et al. The clinicopathologic spectrum of focal cortical dysplasias: A consensus classification proposed by an ad hoc Task Force of the ILAE Diagnostic Methods Commission. Epilepsia. 2011;52(1):158–174. doi:10.1111/j.1528-1167.2010.02777.x

4. Najm I, Lal D, Alonso Vanegas M, et al. The ILAE consensus classification of focal cortical dysplasia: An update proposed by an ad hoc task force of the ILAE diagnostic methods commission. Epilepsia. 2022;63(8):1899–1919. doi:10.1111/epi.17301

5. Hong SJ, Bernhardt BC, Gill RS, Bernasconi N, Bernasconi A. The spectrum of structural and functional network alterations in malformations of cortical development. Brain. 2017;140(8):2133–2143. doi:10.1093/brain/awx145

6. Ripart M, Spitzer H, Williams LZJ, et al. Detection of epileptogenic focal cortical dysplasia using graph neural networks: A MELD study. JAMA Neurol. doi:10.1001/jamaneurol.2024.5406

7. Mellerio C, Labeyrie MA, Chassoux F, et al. Optimizing MR imaging detection of type 2 focal cortical dysplasia: Best criteria for clinical practice. Am J Neuroradiol. 2012;33(10):1932–1938. doi:10.3174/ajnr.A3081

8. Jayalakshmi S, Vooturi S, Vadapalli R, Madigubba S, Panigrahi M. Predictors of surgical outcome in focal cortical dysplasia and its subtypes. J Neurosurg. 2021;136(2):512–522. doi:10.3171/2020.12.JNS203385

9. Colliot O, Antel SB, Naessens VB, Bernasconi N, Bernasconi A. In vivo profiling of focal cortical dysplasia on high-resolution MRI with computational models. Epilepsia. 2006;47(1):134–142. doi:10.1111/j.1528-1167.2006.00379.x

10. Adler S, Lorio S, Jacques TS, et al. Towards in vivo focal cortical dysplasia phenotyping using quantitative MRI. Neuroimage Clin. 2017;15:95–105. doi:10.1016/j.nicl.2017.04.017

11. Hong SJ, Bernhardt BC, Caldairou B, et al. Multimodal MRI profiling of focal cortical dysplasia type II. Neurology. 2017;88(8):734–742. doi:10.1212/WNL.0000000000003632

12. Lorio S, Adler S, Gunny R, et al. MRI profiling of focal cortical dysplasia using multi-compartment diffusion models. Epilepsia. 2020;61(3):433–444. doi:10.1111/epi.16451

13. Snell A, Du J, Vegh V, Reutens D. MRI techniques for detecting focal cortical dysplasia: A systematic review. Seizure. 2026;135:56–72. doi:10.1016/j.seizure.2026.01.003

14. Kala D, Abragimovich T, Janča R, et al. Beyond structural MRI: Combined diffusion metric improve delineation of epileptogenic tissue in focal cortical dysplasia. Sci Rep. doi:10.1038/s41598-026-57292-w

15. Paquola C, Bethlehem RA, Seidlitz J, et al. Shifts in myeloarchitecture characterise adolescent development of cortical gradients. Elife. 2019;8:e50482. doi:10.7554/eLife.50482

16. Paquola C, Seidlitz J, Benkarim O, et al. A multi-scale cortical wiring space links cellular architecture and functional dynamics in the human brain. PLoS Biol. 2020;18(11):e3000979. doi:10.1371/journal.pbio.3000979

17. Paquola C, Royer J, Tsigaras T, et al. Intracortical microstructure profiling: A cross-modal method for indexing cortical lamination. Imaging Neurosci. 2026;4:IMAG.a.1212. doi:10.1162/IMAG.a.1212

18. Hong SJ, Kim H, Schrader D, Bernasconi N, Bernhardt BC, Bernasconi A. Automated detection of cortical dysplasia type II in MRI-negative epilepsy. Neurology. 2014;83(1):48–55. doi:10.1212/WNL.0000000000000543

19. Spitzer H, Ripart M, Whitaker K, et al. Interpretable surface-based detection of focal cortical dysplasias: a Multi-centre Epilepsy Lesion Detection study. Brain. 2022;145(11):3859–3871. doi:10.1093/brain/awac224

20. Vorndran J, Blümcke I. A deep-learning-based histopathology classifier for focal cortical dysplasia (FCD) unravels a complex scenario of comorbid FCD subtypes. Epilepsia. 2024;65(12):3501–3512. doi:10.1111/epi.18161

21. Chen J, Sahlas E, Zhou Y, et al. Use of artificial intelligence in magnetic resonance imaging across the epileptic patient’s journey: A meta-analysis of four clinical applications. Epilepsia. doi:10.1002/epi.70254

22. Hong SJ, Lee HM, Gill R, et al. A connectome-based mechanistic model of focal cortical dysplasia. Brain. 2019;142(3):688–699. doi:10.1093/brain/awz009

23. Sydnor VJ, Larsen B, Seidlitz J, et al. Intrinsic activity development unfolds along a sensorimotor-association cortical axis in youth. Nat Neurosci. 2023;26(4):638–649. doi:10.1038/s41593-023-01282-y

24. Sydnor VJ, Larsen B, Bassett DS, et al. Neurodevelopment of the association cortices: Patterns, mechanisms, and implications for psychopathology. Neuron. 2021;109(18):2820–2846. doi:10.1016/j.neuron.2021.06.016

25. Lee HM, Hong SJ, Gill R, et al. Multimodal mapping of regional brain vulnerability to focal cortical dysplasia. Brain. 2023;146(8):3404–3415. doi:10.1093/brain/awad060

26. Lee HM, Gill RS, Fadaie F, et al. Unsupervised machine learning reveals lesional variability in focal cortical dysplasia at mesoscopic scale. Neuroimage Clin. 2020;28:102438. doi:10.1016/j.nicl.2020.102438

27. Schuch F, Walger L, Schmitz M, et al. An open presurgery MRI dataset of people with epilepsy and focal cortical dysplasia type II. Sci Data. 2023;10(1):475. doi:10.1038/s41597-023-02386-7

28. Bernasconi A, Cendes F, Theodore WH, et al. Recommendations for the use of structural magnetic resonance imaging in the care of patients with epilepsy: A consensus report from the International League Against Epilepsy Neuroimaging Task Force. Epilepsia. 2019;60(6):1054–1068. doi:10.1111/epi.15612

29. Cruces RR, Royer J, Herholz P, et al. Micapipe: A pipeline for multimodal neuroimaging and connectome analysis. Neuroimage. 2022;263:119612. doi:10.1016/j.neuroimage.2022.119612

30. Dale AM, Fischl B, Sereno MI. Cortical surface-based analysis. I. Segmentation and surface reconstruction. Neuroimage. 1999;9(2):179-194. doi:10.1006/nimg.1998.0395

31. Glasser MF, Sotiropoulos SN, Wilson JA, et al. The minimal preprocessing pipelines for the Human Connectome Project. Neuroimage. 2013;80:105–124. doi:10.1016/j.neuroimage.2013.04.127

32. Cappelle S, Pareto D, Sunaert S, et al. T1w/FLAIR ratio standardization as a myelin marker in MS patients. Neuroimage Clin. 2022;36:103248. doi:10.1016/j.nicl.2022.103248

33. Xie K, Sahlas E, Ngo A, et al. Personalized biomarkers of multiscale functional alterations in temporal lobe epilepsy. Nat Commun. 2025;16(1):10145. doi:10.1038/s41467-025-65042-1

34. Jiang L, Zuo XN. Regional homogeneity: A multimodal, multiscale neuroimaging marker of the human connectome. Neuroscientist. 2016;22(5):486–505. doi:10.1177/1073858415595004

35. Cole MW, Pathak S, Schneider W. Identifying the brain’s most globally connected regions. Neuroimage. 2010;49(4):3132–3148. doi:10.1016/j.neuroimage.2009.11.001

36. Fortin JP, Cullen N, Sheline YI, et al. Harmonization of cortical thickness measurements across scanners and sites. Neuroimage. 2018;167:104–120. doi:10.1016/j.neuroimage.2017.11.024

37. Chen J, Ngo A, Rodriguez-Cruces R, et al. Changes in gray matter morphology and white matter microstructure across the adult lifespan in people with temporal lobe epilepsy. Neurology. 2025;105(6):e213688. doi:10.1212/WNL.0000000000213688

38. Vos de Wael R, Benkarim O, Paquola C, et al. BrainSpace: a toolbox for the analysis of macroscale gradients in neuroimaging and connectomics datasets. Commun Biol. 2020;3(1):103. doi:10.1038/s42003-020-0794-7

39. Van Essen DC, Smith SM, Barch DM, et al. The WU-Minn Human Connectome Project: an overview. Neuroimage. 2013;80:62–79. doi:10.1016/j.neuroimage.2013.05.041

40. Feinberg DA, Moeller S, Smith SM, et al. Multiplexed echo planar imaging for sub-second whole brain fMRI and fast diffusion imaging. PLoS One. 2010;5(12):e15710. doi:10.1371/journal.pone.0015710

41. Nakagawa JM, Donkels C, Fauser S, et al. Characterization of focal cortical dysplasia with balloon cells by layer-specific markers: Evidence for differential vulnerability of interneurons. Epilepsia. 2017;58(4):635–645. doi:10.1111/epi.13690

42. Blümcke I, Vinters HV, Armstrong D, Aronica E, Thom M, Spreafico R. Malformations of cortical development and epilepsies: Neuropathological findings with emphasis on focal cortical dysplasia. Epileptic Disord. 2009;11(3):181–193. doi:10.1684/epd.2009.0261

43. Mackay MT, Becker LE, Chuang SH, et al. Malformations of cortical development with balloon cells: Clinical and radiologic correlates. Neurology. 2003;60(4):580–587. doi:10.1212/01.wnl.0000044053.09023.91

44. Bernhardt BC, Hong SJ, Bernasconi A, Bernasconi N. Magnetic resonance imaging pattern learning in temporal lobe epilepsy: Classification and prognostics. Ann Neurol. 2015;77(3):436–446. doi:10.1002/ana.24341

45. Sinha N, Wang Y, Moreira da Silva N, et al. Structural brain network abnormalities and the probability of seizure recurrence after epilepsy surgery. Neurology. 2021;96(5):e758–e771. doi:10.1212/WNL.0000000000011315

46. Huppertz HJ, Grimm C, Fauser S, et al. Enhanced visualization of blurred gray-white matter junctions in focal cortical dysplasia by voxel-based 3D MRI analysis. Epilepsy Res. 2005;67(1-2):35–50. doi:10.1016/j.eplepsyres.2005.07.009

47. Adler S, D’Arco F, Mankad K, Kyncl M, Arzimanoglou A, Marusic P. Harmonization of MRI sequences across ERN EpiCARE centers. Epilepsia Open. 2025;10(2):587–592. doi:10.1002/epi4.13115

48. Yang H, Wu G, Li Y, et al. Connectional axis of individual functional variability: Patterns, structural correlates, and relevance for development and cognition. Proc Natl Acad Sci. 2025;122(12):e2420228122. doi:10.1073/pnas.2420228122

49. Zhang S, Larsen B, Sydnor VJ, et al. In vivo whole-cortex marker of excitation-inhibition ratio indexes cortical maturation and cognitive ability in youth. Proc Natl Acad Sci. 2024;121(23):e2318641121. doi:10.1073/pnas.2318641121

50. Cohen NT, You X, Krishnamurthy M, et al. Networks underlie temporal onset of dysplasia-related epilepsy: A MELD study. Ann Neurol. 2022;92(3):503–511. doi:10.1002/ana.26442

