## Supplemental Material for "Multiscale alterations and cortex-wide organizational trends in focal cortical dysplasia"

*MRI data acquisition*

For the MICs dataset, MRI data were acquired using a 3T Siemens Magnetom Prisma Fit scanner with a 64-channel head coil receiver. We acquired T1w scans via a three-dimensional magnetization-prepared rapid gradient-echo (3D-MPRAGE) sequence (repetition time (TR) = 2300 ms, echo time (TE) = 3.14 ms, inversion time (TI) = 900 ms, flip angle (FA) = 9º, matrix = 320 × 320, voxel size = 0.8 × 0.8 × 0.8 $mm^3$, field of view (FOV) = 256 × 256 $mm^2$, 224 sagittal slices). A three-dimensional FLAIR sequence was also acquired (TR = 5000 ms, TE = 393 ms, TI = 1600 ms, matrix = 256 × 256, voxel size = 1 × 1 × 1 $mm^3$, FOV = 256 × 256 $mm^2$, 192 sagittal slices). We acquired rsfMRI using a two-dimensional blood-oxygen-level-dependent (2D-BOLD) echo-planar sequence with multi-band acceleration (TR = 600 ms, TE = 30 ms, FA = 50º, matrix = 80 × 80, voxel size = 3 × 3 × 3 $mm^3$, FOV = 240 × 240 $mm^2$, multi-band factor = 6, echo spacing = 0.54 ms, 48 slices oriented to the bicommissural line minus 30 degrees). For the ~7-minute duration of the rsfMRI scan, participants fixated a cross displayed in the centre of the screen and were instructed to not fall asleep. For the NOEL and Bonn datasets, details of MRI data acquisition parameters are described elsewhere[26,27,28].

*Correlations across features*

To assess the extent to which different features may vary in tandem within FCD lesions, we computed the Pearson correlation coefficient between each pair of mean within-lesion *w*-scores across patients. CT correlated positively with BL ($r = 0.434$, $p_{FDR} = 1.439*10^{-2}$) and negatively with RH ($r = 0.373$, $p_{FDR} = 4.376*10^{-2}$). BL correlated negatively with M1 ($r = -0.489$, $p_{FDR} = 4.299*10^{-3}$) and M2 ($r = -0.360$, $p_{FDR} = 4.663*10^{-2}$), and positively with M3 ($r = 0.487$, $p_{FDR} = 4.299*10^{-3}$). The two functional features, RH and NS, were also positively correlated ($r = 0.420$, $p_{FDR} = 1.808*10^{-2}$). Interestingly, M4 did not correlate significantly with BL ($r = 0.099$, $p_{FDR} = 6.999*10^{-1}$), suggesting that M4 could carry information independent from BL in differentiating FCD Type IIa and Type IIb.
